# Inflammatory Oxidative Stress Compounds Inhibit Insulin Secretion through Rapid Protein Carbonylation

**DOI:** 10.1101/2025.07.23.666010

**Authors:** Emma F. Saunders, Katherine R. Schultz, Isaiah Lowe, Aimee L. Anderson, Vrishank S. Bikkumalla, David Soto, Nhi Y. Tran, Sharon Baumel-Alterzon, Jefferson D. Knight, Colin T. Shearn

**Affiliations:** Department of Chemistry, University of Colorado Denver, USA; Department of Pediatrics, University of Colorado Anschutz Medical Campus, Aurora, USA; Department of Molecular and Cellular Endocrinology, Arthur Riggs Diabetes & Metabolism Research Institute at City of Hope, Duarte, CA, USA

**Author notes:** Equal contributions (Emma F. Saunders, Katherine R. Schultz).

**Keywords:** Insulin, secretion, carbonylation, inflammation, lysine reactivity, cysteine reactivity, type 1 diabetes, TIRFM, GRINCH, MIN6, INS-1, NOD, reactive aldehyde, pro-inflammatory cytokine, secretory pathway, vesicle trafficking

## Abstract

Pancreatic β-cells in pre-type 1 diabetes (T1D) experience stress due to islet inflammation, which accompanies early defects in insulin secretion that precede autoimmune destruction. One product of inflammatory stress is protein carbonylation (PC), brought on by reactive oxygen species (ROS) combining with lipids to produce reactive aldehydes such as 4-hydroxynonenal (4-HNE) that irreversibly modify Cys, His, and Lys sidechains. In this study, we used proteomics to measure patterns of PC in pancreatic islets from 10-week-old pre-diabetic NOD mice and in cultured insulin-secreting cells treated with either 4-HNE or pro-inflammatory cytokines. All three stress conditions increased carbonylation of proteins central to β-cell function including Rab GTPases and other proteins that are essential for vesicle trafficking. Gene ontology analysis indicates that the affected proteins and pathways in pre-diabetic NOD islets reflect a combination of those impacted by 4-HNE and cytokine treatment. Furthermore, both 4-HNE and cytokines significantly inhibited insulin secretion by ∼50% in cultured MIN6 and INS-1-GRINCH cells. In particular, exposure to 4-HNE for as little as 5 minutes suppressed insulin secretion and increased the carbonylation of over 1000 proteins. Overall, the observed PC pattern in pre-T1D islets is consistent with a model in which β-cells experience multiple sources of oxidative stress, including ROS generation within β-cells themselves and reactive compounds released by infiltrating immune cells. The latter exogenous source may represent a novel rapid mechanism for inhibiting insulin secretion.

## Introduction

With over 1.8 million cases diagnosed in the U.S. in recent years, type 1 diabetes (T1D) remains a major public health concern (1; 2). T1D is brought on via environmental stressors such as viral infections which initiate an infiltration of autoimmune cells, including T cells and macrophages, into pancreatic islets (3–7). Effects of the immune infiltration on β-cells ultimately lead to β-cell destruction (1; 8). Intriguingly, studies report a progressive decline in β-cell secretory function prior to cell death during the development of T1D and that patients with preserved β-cell function exhibit a better response to insulin therapy, suggesting that improving β-cell function may be a useful additional strategy to delay the onset of T1D (9; 10). However, the molecular mechanisms that provoke β-cell dysfunction prior to the onset of T1D remain underexplored.

Growing evidence suggests that oxidative stress, characterized by increased production of reactive oxygen species (ROS), plays a significant role in the pathology of T1D (11). This is not surprising, given that β-cells express low levels of key antioxidant enzymes, making them highly susceptible to oxidative damage (12–15). While a basal level of ROS production in healthy mitochondria is important for maintaining cellular homeostasis, β-cells can be overwhelmed during insulitis by additional ROS production that arises from at least two sources (16). First, endogenous β-cell ROS production and leakage increase as a result of ER stress (17; 18) and from proinflammatory cytokines such as IL-1β, IFNγ and TNFα, which activate β-cell signaling pathways that regulate mitochondrial function (11; 19-26). Second, pro-inflammatory macrophages activate NADPH oxidase 2 (NOX2) to produce and release superoxide anions and other ROS independent of mitochondria (4; 27). ROS can readily penetrate β-cell plasma and subcellular membranes, where they react with polyunsaturated fatty acid chains to produce oxidized lipid products such as reactive aldehydes, which diffuse locally and react with proteins (28). Reactive lipid aldehydes can also travel further from the site of production, as evidenced by their detection in serum of diabetic patients (29; 30).

Recent studies have emphasized the roles of oxidative stress-derived post-translational protein modifications in the development of T1D (31; 32). Among these is protein carbonylation (PC), a modification that occurs spontaneously when lipid aldehydes react with cysteine, lysine, and histidine residues (33; 34). The resulting covalent modifications are not enzymatically reversible and can result in protein inactivation, misfolding, aggregation, degradation, and/or the generation of neo-epitopes (35; 36). The accumulation of reactive lipid aldehydes, including 4-hydroxynonenal (4-HNE), has been linked to various pathophysiological conditions, including diabetes and inflammation (37–41); 4-HNE has also been shown to suppress insulin secretion in isolated rat islets (42). Notably, a recent study identified a few dozen carbonylated proteins in islets from pre-diabetic NOD mice (36). Yet, the full extent of inflammation-derived PC in β-cells and its contribution to impaired insulin secretion are not understood.

We recently reported the use of a precise and sensitive method for the global identification of cellular PC. This method, which is based on biotin hydrazide derivatization combined with global LC-MS/MS, enabled the identification of hundreds of carbonylated proteins in liver tissue isolated from human patients diagnosed with metabolic fatty liver disease (43). In the present study, we applied a similar approach to investigate how PC affects β-cell function under conditions associated with T1D. For this purpose, a global PC analysis was performed in pre-diabetic 10-week-old female NOD mice undergoing islet inflammation and in cultured MIN6 β-cells treated with the reactive lipid aldehyde 4-HNE or pro-inflammatory cytokines. To better understand how inflammation-mediated PC affects β-cell function, global PC profiles were correlated with measurements of insulin secretion in MIN6 and INS-1 (GRINCH) β-cells. Overall, we identify hundreds to thousands of carbonylated proteins in pre-diabetic NOD mice and in cell culture conditions approximating pre-T1D inflammatory and oxidative stress. These proteins include several Rab isoforms and other secretory pathway proteins. Both cytokines and reactive aldehydes inhibit insulin secretion by ∼50%, the latter in only a few minutes, consistent with proteomic data showing that PC directly impacts key proteins in the secretory pathway. To our knowledge, this is the first proteome-scale study of protein carbonylation in β-cells and its correlation to impaired insulin secretion.

## Research Design and Methods

### Reagents

4-HNE in ethanol (Cayman Chemical) was evaporated under nitrogen and dissolved in 100% acetonitrile to a concentration of 64 mM and stored under desiccation at -20 °C prior to use. Cytokines IL-1β and IFNγ were from R&D Systems; TNFα was from BD Biosciences. Cytokines were diluted in sterile phosphate-buffered saline (PBS) to the desired stock concentrations (IL-1β 5 µg/mL, IFNγ 100 µg/mL, and TNFα 50 µg/mL), and flash-frozen aliquots were stored at -80°C. Cell culture media, HEPES, sodium pyruvate, β-mercaptoethanol, penicillin/streptomycin, and trypsin were from Gibco. Fetal Bovine Serum (FBS) was from Atlanta Biologicals. Neomycin, Triton X-100 and D-glucose were from ThermoFisher.

### Isolation of murine pancreatic islets

Isolation of mouse islets was conducted under a protocol approved by the University of Colorado Anschutz Medical Campus Institutional Animal Care and Use Committee as described previously (44). Ten-week-old female BALB/c mice (control) or nondiabetic NOD mice were anesthetized and the pancreas was inflated via the common bile duct with collagenase solution and then removed. Following incubation at 37 °C to facilitate digestion, islets were isolated by density centrifugation. Islets were handpicked under a microscope and aliquoted into microcentrifuge tubes containing medium supplemented with fetal bovine serum (5%). Tubes were spun at 300g for 5 min, supernatant was removed, and the tubes were placed in liquid nitrogen to rapidly freeze the islet pellets. Islets were stored at -80 °C until use. Because individual mice only yield approximately 150–250 islets, islets were harvested from 10 mice of the same strain at a time and islets from all of the mice were pooled.

### Cell Culture

INS-1 (GRINCH) cells (45) were cultured in RPMI 1640 media with 10% FBS, 1% penicillin/streptomycin, 1 mM sodium pyruvate, 10 mM HEPES buffer, and 50 µM β-mercaptoethanol. MIN6 cells were cultured in DMEM media with 10% FBS, 1% penicillin/streptomycin, and 50 µM β-mercaptoethanol with or without 0.5% neomycin. Both cell lines were maintained at 40-80% confluency at 37 !ZC with 5% CO₂.

### Insulin ELISA

Approximately 4 x 10^5^ cells were plated in wells on a 24-well plate and allowed to adhere overnight prior to treatment. For 4-HNE treatment, cells were first incubated overnight in assay buffer (114 mM NaCl, 4.7 mM KCl, 1.2 mM KH_2_PO_4_, 1.16 mM MgSO_4_, 40 mM HEPES, 2.5 mM CaCl_2_, 25.5 mM NaHCO_3_ 2% Bovine Serum Albumin (BSA)) containing 2.8 mM glucose, then glucose-starved in assay buffer with 2.8 mM glucose for 1-2 hr prior to addition of the indicated concentration of 4-HNE in BSA-free assay buffer containing 2.8 mM glucose. For cytokine treatment, cells were cultured for the indicated time at 37 !ZC and 5% CO₂ with 10 ng/mL TNFα, 100 ng/mL IFNγ and 5 ng/mL IL-1β in culture media lacking β-mercaptoethanol, then glucose starved in assay buffer with 2.8 mM glucose for 1 h. Cells were then placed into assay buffer with 2.8 mM glucose (control) or the indicated stimulation condition for 1 hour.

To prepare secreted insulin samples, 600 μL from each well of the 24-well plate was centrifuged at 17,000 *× g* for 5 minutes, then added to 500 μL PBS/1% BSA/0.02% sodium azide. To prepare insulin content samples, cells were scraped and lysed by trituration in 0.1% Triton X-100 after secretion samples were removed. Each lysate (∼600 μL) was added to 500 μL PBS/1% BSA/0.02% sodium azide.

Content and secretion samples were diluted into 0.2% BSA/PBS using dilution factors estimated empirically based on DNA concentration (Quant-iT PicoGreen dsDNA assay kit, Thermo Fisher). Insulin ELISAs were conducted using the Mouse Ultrasensitive Insulin ELISA kit (ALPCO) with an overnight incubation at 4°C. Data were analyzed using a five-parameter logistic fit to determine content and secreted insulin concentrations.

### Cell Death Measurement

Approximately 4 x 10^5^ cells were plated in wells on a 24-well plate and were treated as described above. DAPI was added to each well to a concentration of 1 μg/ml and cells were imaged using a Cytation 3 plate reader (BioTek, Winooski, WI). Cell death was quantified by counting DAPI-positive cells before and after addition of 0.015% Triton X-100, which renders all cells DAPI-permeable. Data shown are average ± SEM of 3-6 images each from 3-4 biological replicates.

### Cell Plating and Treatment for TIRFM

Cells were plated at a density of ∼7 × 10^5^ per cm^2^ onto 35-mm glass bottom petri dishes (World Precision Instruments or MatTek) coated with poly-D-lysine. Cells were incubated overnight at 37 !ZC with 5% CO₂ to allow the cells to adhere to the glass, then media was exchanged, and cells were allowed to rest overnight. Cytokine samples were treated for 72 h. Prior to imaging, cells were glucose- and serum-starved in the appropriate media for 30 min and then exchanged into imaging buffer (32 mM HEPES, 145 mM NaCl, 5.6 mM KCl, 0.5 mM MgCl_2_, 2 mM CaCl_2_, 3 mM glucose).

### Microscopy

Cells were imaged using a Zeiss Axioskop microscope equipped with a 488 nm laser. A microperfusion system from ALA Scientific was used to stimulate individual groups of a small number of cells, allowing for ∼8-10 recordings of stimulated secretion to be obtained from different areas of the same dish. For each selected area, a video was recorded with imaging buffer flowing through the perfusion needle for 10 s, followed by 60 s flowing stimulation solution (32 mM HEPES, 80 mM NaCl, 60 mM KCl, 5 mM CaCl_2_, 3 mM glucose).

For experiments studying 4-HNE effects, ∼3 areas per dish were imaged as described above prior to 4-HNE addition. Then, 100 μM 4-HNE (or vehicle) was added and additional areas were imaged over a total time of approximately 15 minutes thereafter. The frequency of secretion events from cells stimulated prior to 4-HNE or vehicle addition was statistically indistinguishable from that after vehicle addition (**Supporting Information, Fig. S1**).

### Microscopy Data Analysis

Videos of cell secretion were analyzed using Fiji/ImageJ. The number of cells and area of cells visible in each video, as well as the number of visible vesicles and number of secretion events in each cell, were counted using the Time Series Analyzer V3 plugin.

### Carbonyl proteomics

For murine islet experiments, ∼1750 islets were pooled from 10 WT (BALB/c) or 10 NOD mice and used for proteomic analysis. For cell culture experiments, ∼1.8 × 10^7^ cells per treatment group were used for proteomic analysis, with 3 replicate treatment groups at each condition. For 4-HNE experiments, 100 µM 4-HNE or vehicle was added to cells in serum-free and glucose-free culture media. For cytokine treatment, cells were cultured with media lacking β-mercaptoethanol and including 10 ng/mL TNFα, 100 ng/mL IFNγ, and 5 ng/mL IL-1β at 37 °C and 5% CO_2_ for the designated amount of time (including 1 h for the 4-HNE control samples and 72h for the cytokine control samples), then scraped and centrifuged at 500 × g for 5 min. Pellets were washed with HBSS, flash frozen in liquid nitrogen, and stored at -80 !ZC.

Cells were lysed by probe sonication (VibraSonics, 3 mm tip) for 3×10 s in lysis buffer (1 mM NaCl, 0.05 mM EDTA, 1 µg/mL antipain, 1 µg/mL leupeptin, 1 µg/mL pepstatin, 1 µg/mL aprotinin, 200 µM PMSF, 50 mM Tris pH 7.5) and centrifuged at 15800 × g for 2 minutes to remove insoluble debris. Total protein content in the supernatant was quantified via BCA assay.

Biotin hydrazide derivatization, purification of aldehyde modified proteins and LC-MS analysis was performed using an Agilent 6550 Quadrupole Time-of-Flight mass spectrometer and Agilent Profinder software as previously described using 1000 μg of protein from each group (46).

### Bioinformatics

The protein lists identified from the mass spectrometry data were analyzed using DAVID (https://david.ncifcrf.gov/summary.jsp). Protein lists (in the **Supporting Information**) were uploaded to DAVID as UniProt IDs and analyzed as a Functional Annotation Chart using either GOTERM_BP_DIRECT (“Biological Process” in data Tables) or GOTERM_CC_DIRECT (“Cellular Compartment” in data Tables) as the Gene Ontology category. The 10 most significant terms are listed in each reported data table. We also analyzed the lists using UP_KW_PTM and GOTERM_MF_DIRECT terms but found these post-translational modification and molecular function analyses were less informative. The most significant enrichment trends were that carbonylation affects proteins that can be acetylated (which makes chemical sense because carbonylation targets exposed lysine residues) and nucleotide binding proteins (including many GTPases as noted in the Results).

Venn diagrams were generated from the protein lists in the **Supporting Information** using the Gent University Bioinformatics toolset (http://bioinformatics.psb.ugent.be).

### Treatment of Cells with Pro-Inflammatory Cytokines for mRNA Analysis and Griess assay

Cells were grown to ∼70% confluency in 6-well plates (∼6 × 10^5^ cells per well) in the appropriate media lacking β-mercaptoethanol. All cells were incubated at 37 °C and 5% CO_2_ for the indicated amount of time, with the control cells sitting for 48 hours. The media was removed and flash frozen for analysis via Griess assay. Cells were lysed with TRIzol reagent (Invitrogen) and stored at -20!ZC for qPCR analysis. To determine total nitrites, a Griess reagent kit (Molecular Probes) was used according to the manufacturer’s instructions with absorbance determined at 548nm.

qRT-PCR to examine mRNA expression was performed using TaqMan probes from Applied Biosystems (Foster City, CA) as previously described (47). For quantification, all data was normalized to the reference gene Hypoxanthine guanine phosphoribosyltransferase (Hprt) and then fold change (Relative expression) was calculated using the 2-ΔΔCt method.

### Statistical Analysis

These data are presented as means ± standard error of the mean (SEM). Comparisons between control and cytokine treated cells were accomplished by 1-way analysis of variance. Statistical significance was set at P<0.05. Prism 5 (GraphPad Software, San Diego, CA) was used to perform all statistical tests. Outliers were determined using the ROUT (Q=1%) in PRISM.

## Results

### Islets from pre-diabetic NOD mice exhibit increased PC in proteins associated with vesicle trafficking

At 10 weeks of age, no significant difference in overall body weight was observed between female NOD mice and their control counterparts (**Fig 1A**). However, in agreement with their pre-diabetic phase, a small but significant increase was observed in NOD blood glucose levels (1.3-fold compared to control mice) (**Fig 1B**).

**Figure 1:**
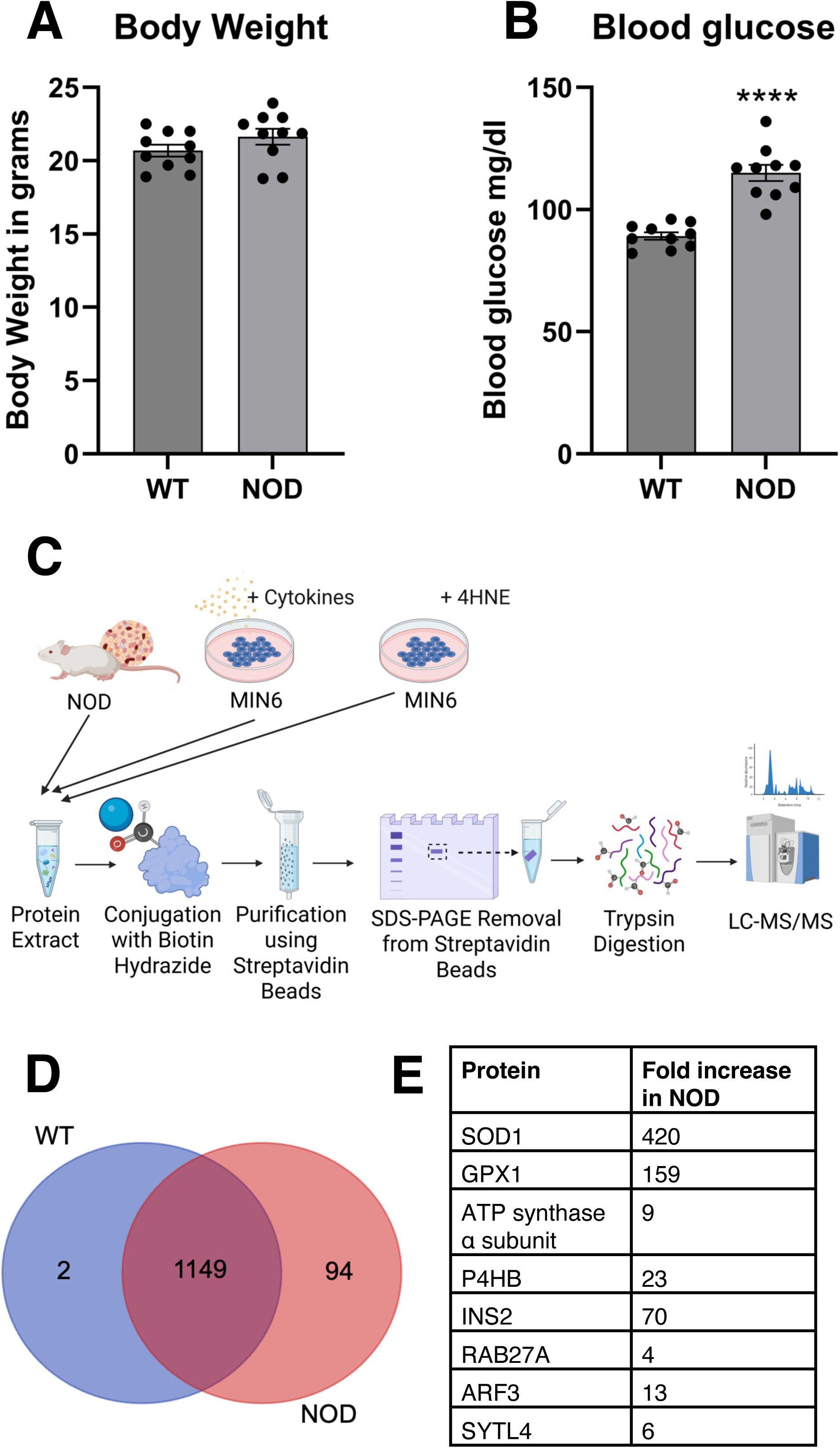
Analysis of islets from pre-diabetic NOD mice. **A:** Body weight and **B:** Blood glucose concentrations at sacrifice in the 10-week-old female NOD mice compared to WT controls. ****p < 0.0001. **C:** Workflow for proteomic analysis of carbonylated proteins. This figure was created using BioRender.com. **D:** Venn diagram of distinct proteins with at least 4 peptides detected by LC-MS following purification of carbonylated proteins from the indicated islet pools. **E:** Selected proteins with increased carbonylation in NOD islets. A full list is in **SI File S1**.

It is estimated that only up to 10% of cellular protein is susceptible to carbonylation under various disease conditions (48). Therefore, to obtain sufficient sample for globally mapping PC, islets from 10 mice were pooled together from NOD or control groups. One mg of total protein from each pooled group was reacted with the carbonyl-specific tag biotin hydrazide, followed by purification using streptavidin beads. Modified proteins were then subjected to in-gel trypsin digestion and analyzed using LC-MS/MS (**Fig. 1C**). A total of 1243 carbonylated proteins were identified from the pooled NOD islet sample (**Fig. 1D**), of which 96% (1196 proteins) were at least 2-fold enriched above control levels based on the MS peak area (**Supporting Information, File S1**).

To identify cellular pathways differentially impacted by carbonylation, these 1196 proteins were subjected to Gene Ontology (GO) enrichment analysis. We observe a statistically significant enrichment of PC in proteins involved in intracellular protein transport, protein folding, translation, and catabolism (**Table 1**). These proteins are correlated with various cellular localizations, including the cytosol, mitochondria, and secretory vesicles (the latter being among the compartments identified as ’synapse’ in our GO analysis), suggesting that PC occurs throughout the cell (**Table 1**). Selected proteins representing these pathways and localizations are listed in **Fig. 1E**. In agreement with previously reported data, protein disulfide isomerase P4Hb (36) and several subunits of ATP synthase (49) were identified as carbonylated in NOD islets. Other carbonylated proteins included the antioxidant proteins glutathione peroxidase (GPX1) and superoxide dismutase (SOD1) as well as the products of the *INS2* gene; i.e., insulin and/or proinsulin (**Fig. 1E**). Interestingly, many proteins involved in vesicle trafficking, exocytosis, and endocytosis were among those with increased PC in the NOD islets, including the COPII coat GTPase Sar1b, Arf GTPases, Rab GTPases, the Rab27a effector protein granuphilin/Slp-4, the secretory vesicle docking protein Munc18-1, N-ethylmaleimide sensitive factor (NSF), and adaptor protein 2 (AP2) complex subunits (**Fig. 1E, Table S1, and SI File S1**).

**Table 1:**
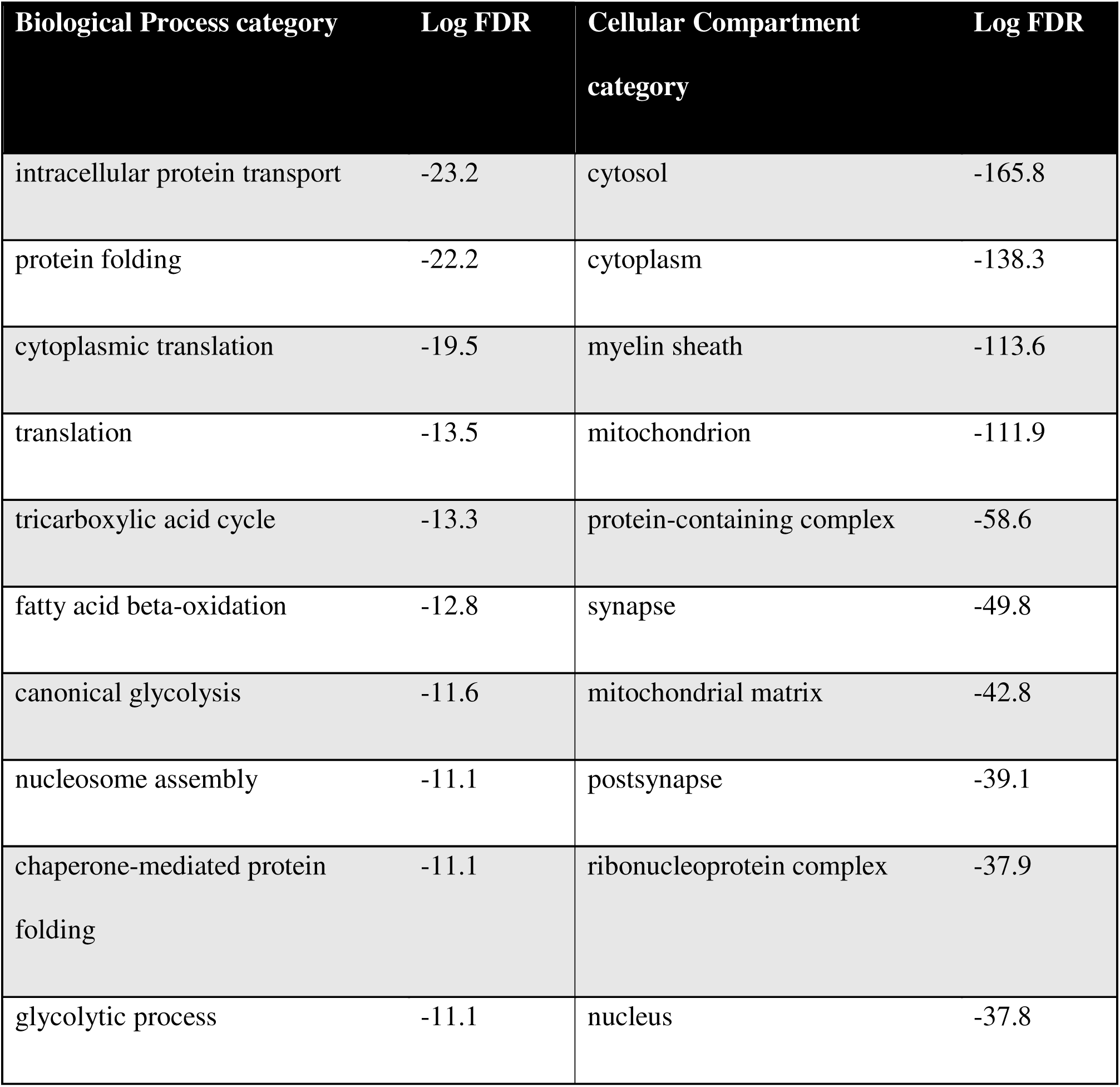
Gene Ontology analysis of carbonylated proteins at least 2-fold enriched in NOD mouse islets.

| Biological Process category | Log FDR | Cellular Compartment category | Log FDR |
| --- | --- | --- | --- |
| intracellular protein transport | -23.2 | cytosol | -165.8 |
| protein folding | -22.2 | cytoplasm | -138.3 |
| cytoplasmic translation | -19.5 | myelin sheath | -113.6 |
| translation | -13.5 | mitochondrion | -111.9 |
| tricarboxylic acid cycle | -13.3 | protein-containing complex | -58.6 |
| fatty acid beta-oxidation | -12.8 | synapse | -49.8 |
| canonical glycolysis | -11.6 | mitochondrial matrix | -42.8 |
| nucleosome assembly | -11.1 | postsynapse | -39.1 |
| chaperone-mediated protein folding | -11.1 | ribonucleoprotein complex | -37.9 |
| glycolytic process | -11.1 | nucleus | -37.8 |

### β-cells exposed to 4-HNE exhibit increased PC and impaired insulin secretion

Because islets from pre-diabetic NOD mice exhibited a significant increase of PC in proteins associated with vesicle trafficking, we decided to test the effect of PC on insulin secretion under stresses associated with inflammation. For this purpose, we used cultured β-cell lines to examine the effects of two physiological sources of carbonylation stress: exogenous reactive lipid aldehydes (approximating ROS release from macrophages, which trigger lipid aldehyde formation in nearby cellular membranes) and proinflammatory cytokines (approximating their production during inflammation).

Previous studies have shown that the reactive lipid aldehyde 4-HNE can suppress insulin secretion (42). To validate this finding, MIN6 β-cells were treated with 4-HNE and glucose-stimulated insulin secretion was measured using ELISA. As expected, insulin secretion was potently inhibited by one-hour treatment with 4-HNE over the physiologically relevant range of 10 to 100 μM (50) (**Fig. 2A**). Surprisingly, treatment with 100 μM 4-HNE for as little as 5 min also significantly inhibited insulin secretion (**Fig 2A**).

**Figure 2:**
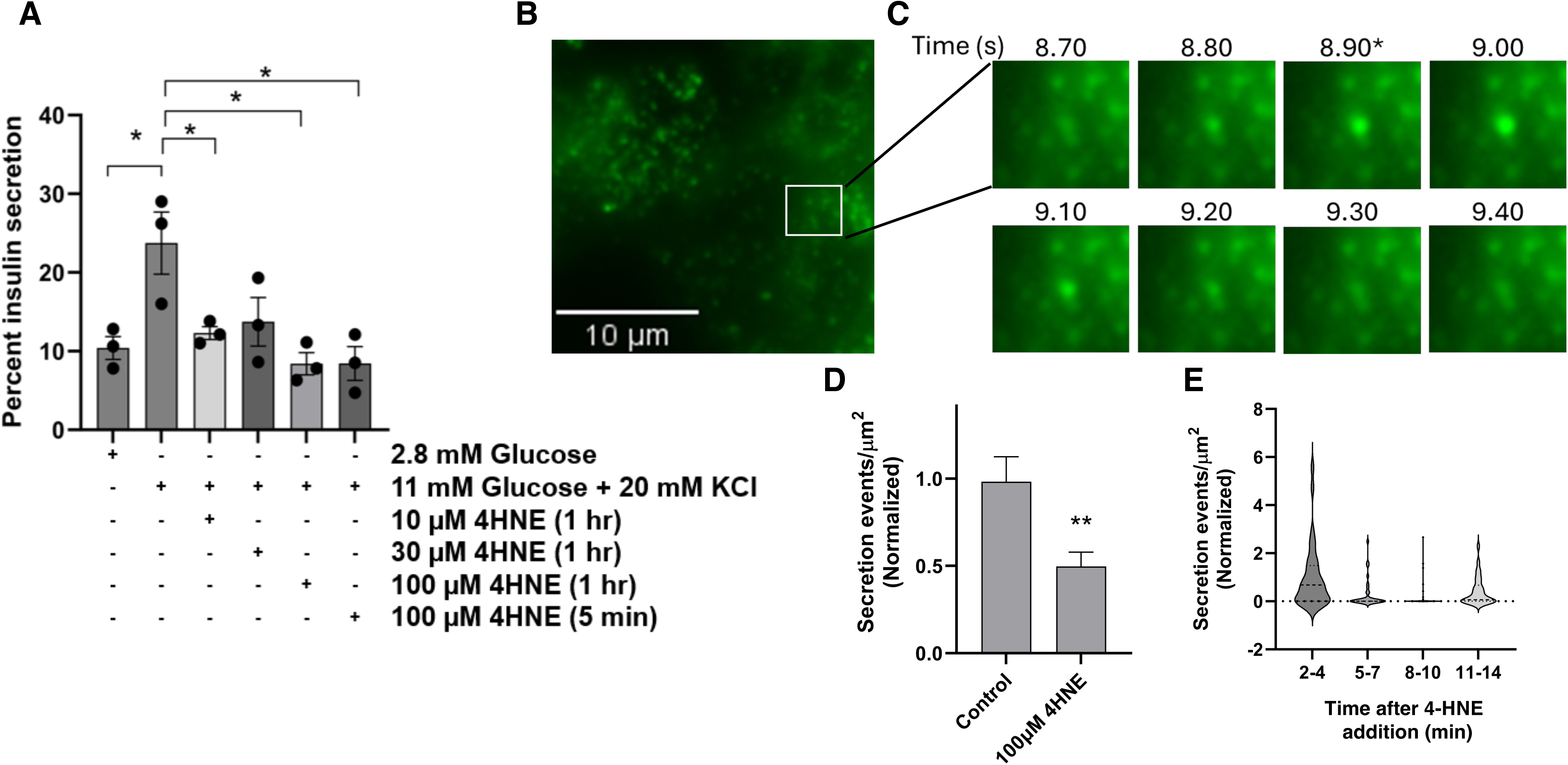
4-HNE inhibition of insulin secretion. **A:** Insulin secretion was measured from MIN6 cells using ELISA after 4-HNE treatment as indicated followed by stimulation with glucose + KCl. *p < 0.05. **B:** Representative image of INS-1 (GRINCH) cells following stimulation by perfusion of 60 mM KCl. Scale bar: 10 μm. **C:** Time series of the region marked by the box in Panel B, illustrating observation of a single secretion event. Frames are stamped with the elapsed time since initiation of KCl perfusion, and the onset of the secretion event is marked with an asterisk. **D:** Quantification of the number of secretion events per minute per square micron of cell surface area among all INS-1 (GRINCH) cells treated with 100 μM 4-HNE as compared to control, normalized to the average of the control cells. **p < 0.01. **E:** Violin plot of the number of secretion events per minute per μm^2^ of INS-1 (GRINCH) cells broken down by the time after 4-HNE addition.

Protein carbonylation reactions typically occur within seconds to minutes of 4-HNE addition (51; 52). To gain improved time resolution and subcellular insight on the effects of 4-HNE on insulin exocytosis, we performed total internal reflection fluorescence microscopy (TIRFM) using the INS-1 (GRINCH) cell line, which expresses sfGFP fused to C-peptide, rendering secretory granules fluorescent (45). Cells were treated with 100 μM 4-HNE or vehicle control and then stimulated via KCl perfusion for 1 minute, during which release of individual insulin granules was imaged using TIRFM (**Fig 2B-C**). Pre-treatment with 4-HNE decreased the number of secretory events by 50% compared to untreated controls (**Fig 2D**), consistent with the GSIS results. This effect was strongest between 5-15 min after 4-HNE addition (**Fig 2E**). Cell death was not increased under these conditions, as measured using trypan blue staining **(Fig. S2**). To identify protein targets of carbonylation by exogenous reactive aldehydes, we performed LC-MS/MS on carbonylated proteins (**Fig. 1C**). MIN6 cells were treated with 100 μM 4-HNE for 5 or 60 minutes. Cells were then lysed and carbonylated proteins were isolated using the biotin hydrazide streptavidin purification method (46) and identified using LC-MS/MS. Overall, 2509 proteins were identified in either the vehicle control and/or the cells treated with 4-HNE for 5 minutes (**Fig 3A and SI File S2**). Of these, 2037 proteins were at least 2-fold enriched in the treated samples relative to controls (**Fig 3B**). In the 60-min 4-HNE treated samples, a total of 2471 proteins were identified, among which 2215 proteins had > 2-fold increased peak area relative to controls (**Fig 3 and SI File S2**). Some proteins, including syntaxin-5 (STX5) and histone H3.1, reached their peak level of carbonylation by 5 min, while most other carbonylated proteins continued to increase in abundance over 60 min (**Fig. 3C**).

**Figure 3:**
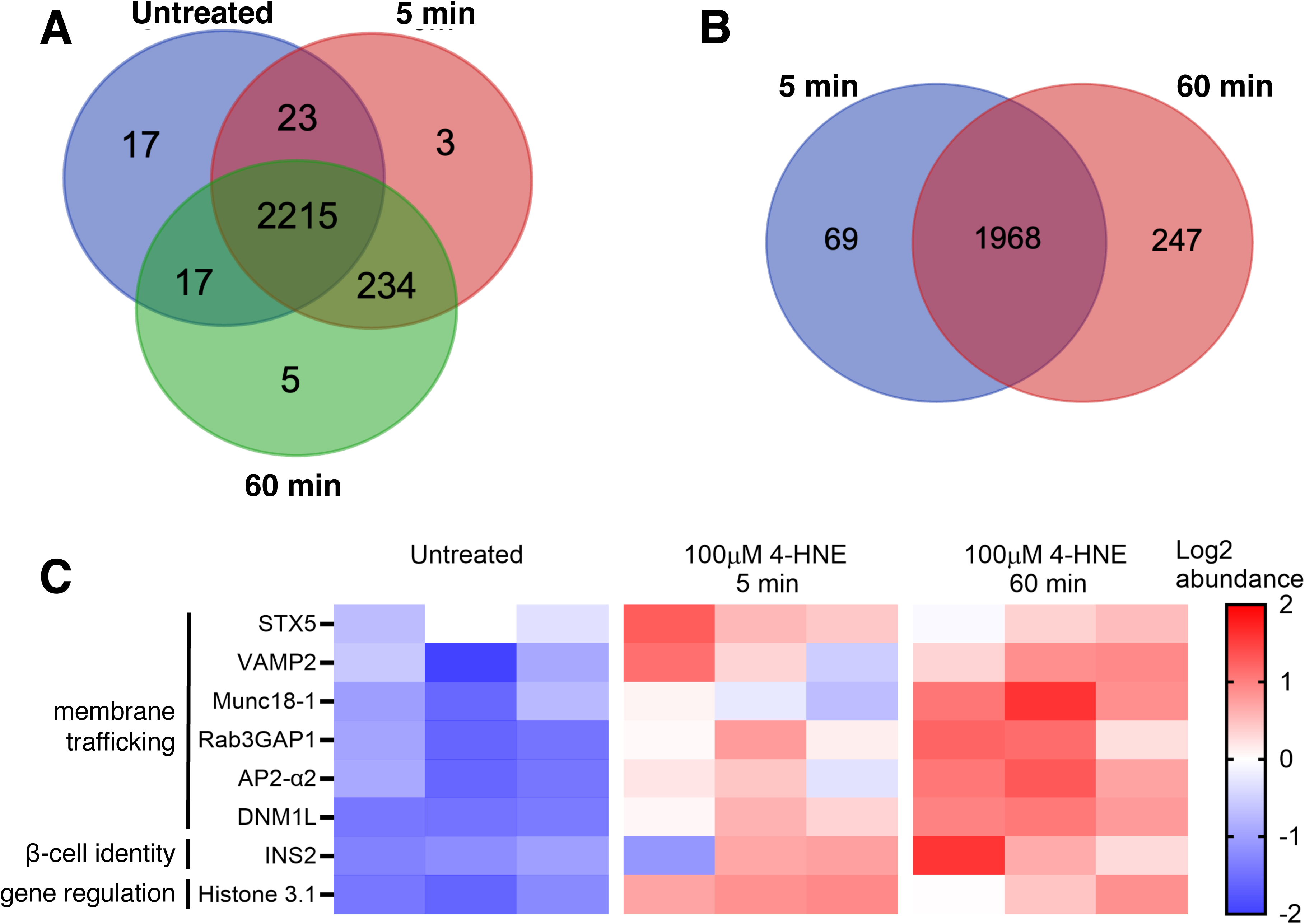
Carbonylated proteins in MIN6 cells treated with 100. μ**M 4-HNE. A:** Venn diagram showing the number of distinct proteins detected by LC-MS following purification of carbonylated proteins from cells treated with 4-HNE or control samples for the indicated time. Only proteins from which four or more distinct peptides were detected are included. **B:** Venn diagram showing the number of distinct proteins for which ≥ 2x greater spectral peak area was observed in the treatment samples relative to controls. **C:** Heat map showing relative abundance of PC for representative selected proteins. Full lists are given in **File S2**.

GO enrichment analysis of the 2215 proteins with > 2-fold increased carbonylation after 60 min 4-HNE treatment revealed several preferentially targeted biological processes in common with the NOD islets, including translation and intracellular protein transport (which includes vesicular transport) (**Table 2**). The most significantly impacted cellular compartments were cytoplasm and nucleus, while synapse (including secretory vesicles) and mitochondria (including ATP synthase) were also among the top 10 impacted categories in this analysis. GO enrichment analysis of the 5-minute treatment yielded similar results (**Table S2**).

**Table 2:** GO categories among carbonylated proteins 2-fold enriched following 60-minute 4-HNE treatment.

| **Biological Process category** | **Log FDR** | **Cellular Compartment category** | **Log FDR** |
| --- | --- | --- | --- |
| cytoplasmic translation | -48.9 | cytosol | -280.7 |
| intracellular protein transport | -47.0 | cytoplasm | -243.5 |
| protein transport | -46.9 | nucleus | -151.7 |
| translation | -39.6 | nucleoplasm | -123.3 |
| mRNA processing | -38.1 | postsynapse | -83.3 |
| translation at postsynapse | -36.9 | protein-containing complex | -76.7 |
| translation at presynapse | -36.0 | ribonucleoprotein complex | -73.6 |
| cell division | -30.9 | synapse | -68.0 |
| chromatin remodeling | -29.0 | mitochondrion | -62.6 |
| protein stabilization | -23.6 | presynapse | -56.2 |

### Effects of proinflammatory cytokines on insulin secretion and protein carbonylation

Proinflammatory cytokines are important contributors to the pathogenesis of T1D and are known to increase ROS production and apoptosis in β-cells (5; 53). To examine the impact of proinflammatory cytokines on oxidative stress in cell culture, MIN6 cells were treated with a cocktail of cytokines (10 ng/mL TNFα, 100 ng/mL IFNγ, and 5 ng/mL IL1β) for 24, 48 and 72 h. Cytokine exposure resulted in a significant increase in nitrite production throughout the time course as measured using a Griess assay (**Fig 4A**). Because nitrite production typically accompanies oxidative stress, we subsequently sought to determine the effects of proinflammatory cytokines on the expression of selected antioxidant enzyme genes Nqo1, Gclc and Sod2 using qPCR (24). Interestingly, expression of all three genes significantly decreased at 24 and 48 h post-cytokine exposure. By 72 h, expression recovered to levels not significantly different compared to untreated cells (**Fig 4B**). These results suggest that these cells require up to 72h to mount an antioxidant response under these conditions, during which time reactive nitrogen species (and therefore ROS) continue to accumulate.

**Figure 4:**
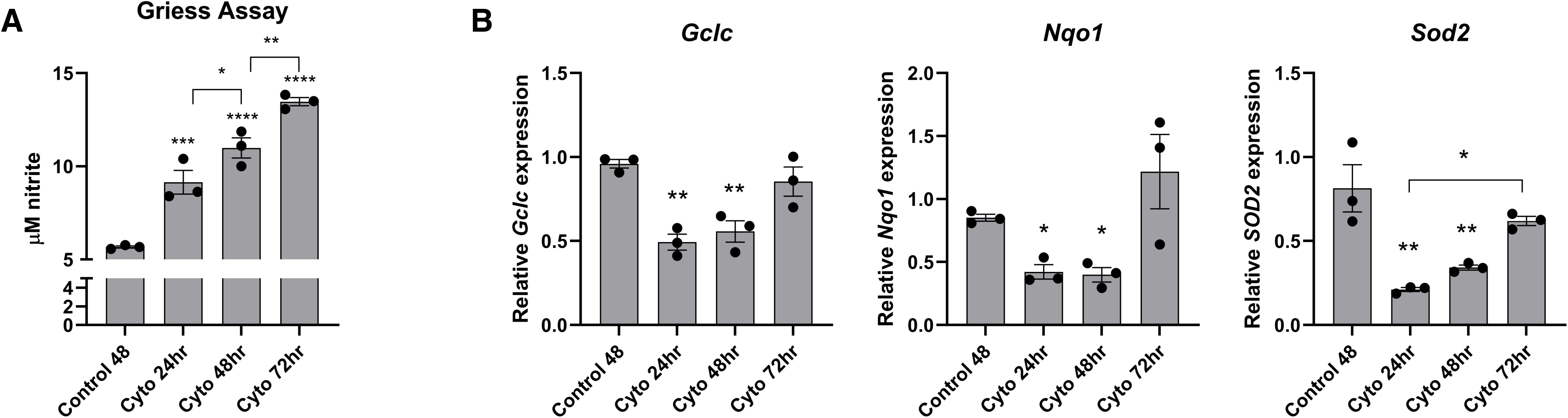
Oxidative and nitrosative stress response following cytokine treatment of MIN6 cells. **A:** Nitrite quantitation from a Griess assay performed on the media removed from the cells following the indicated times of exposure to the cytokine cocktail. **B:** Quantitation of gene expression using qPCR for the oxidative stress reporter genes Gclc, Nqo1 and SOD2. *p < 0.05, **p < 0.01, ***p < 0.001, ****p < 0.0001.

To examine the impact of this cytokine-mediated oxidative stress on insulin secretion, MIN6 β-cells were treated with the proinflammatory cytokine cocktail for 72 h followed by quantification of GSIS. Compared to untreated cells, 72-h cytokine exposure significantly suppressed insulin secretion by over 50% (**Fig 5A**). Similarly, fusion of insulin secretory granules was suppressed by ∼50% in INS-1 (GRINCH) cells following 72-h cytokine exposure as measured using TIRFM (**Fig 5B**). Both of these results are consistent with previous reports that proinflammatory cytokines suppress insulin secretion in INS-1 cells. (54) Trypan blue assays showed a significant increase in cell death after 72-h cytokine treatment, as expected based on previous studies (25) (**Fig. S2**).

**Figure 5:**
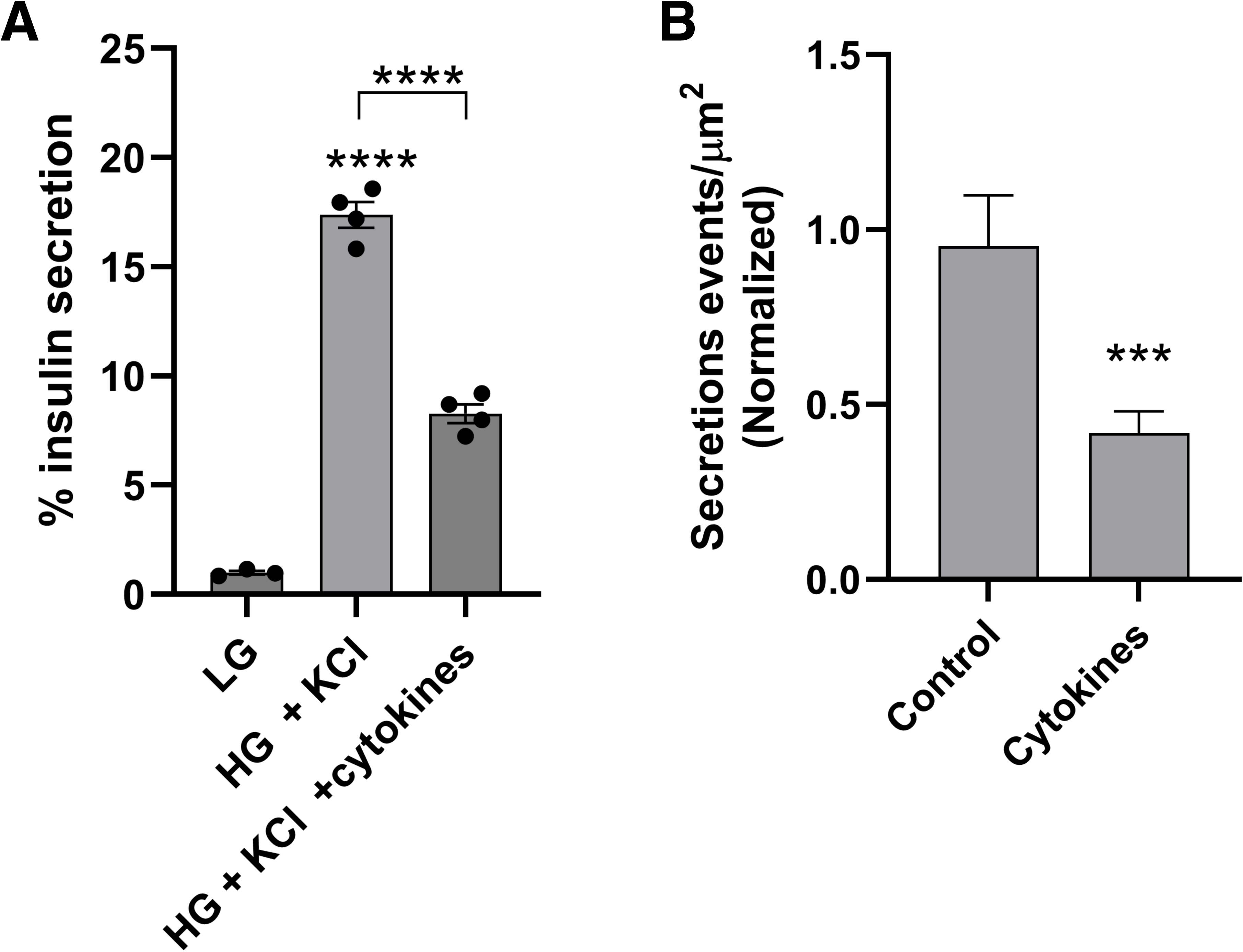
Inhibition of insulin secretion by proinflammatory cytokines. **A:** Insulin secretion was measured from MIN6 cells using ELISA after 72-h cytokine treatment followed by stimulation with 11 mM glucose (HG) and 20 mM KCl, compared to a 2.8 mM glucose (LG) control. **B:** Insulin secretion was measured from INS-1 GRINCH cells using TIRFM following local perfusion of 60 mM KCl onto control vs. 72-h cytokine treated cells. ***p < 0.001; ****p < 0.0001.

Because proinflammatory cytokines are known to increase ROS production (24), we next identified carbonylated proteins from MIN6 cells before and after cytokine treatment in order to correlate secretory dysfunction with PC. Overall, a total of 1761 proteins were identified in samples treated with cytokines for either 36h or 72h or the vehicle control, the large majority of which were identified in all three samples (**Fig 6A**). Due to the longer timescale of the cytokine experiments relative to the 4-HNE experiments, we used a lower fold increase threshold for bioinformatics analysis of these samples, 1.5-fold rather than 2-fold. The 36h cytokine treatment produced only 39 proteins with this level of enrichment, while 72h treatment produced 535 carbonylated proteins with >1.5-fold greater peak area compared to controls (**Fig 6B**). This time lag is consistent with the Griess assay and qPCR results above, which suggest an accumulation of ROS and RNS over 72 h. The abundance of selected carbonylated proteins from these experiments are quantified in **Fig. 6C**.

**Figure 6:**
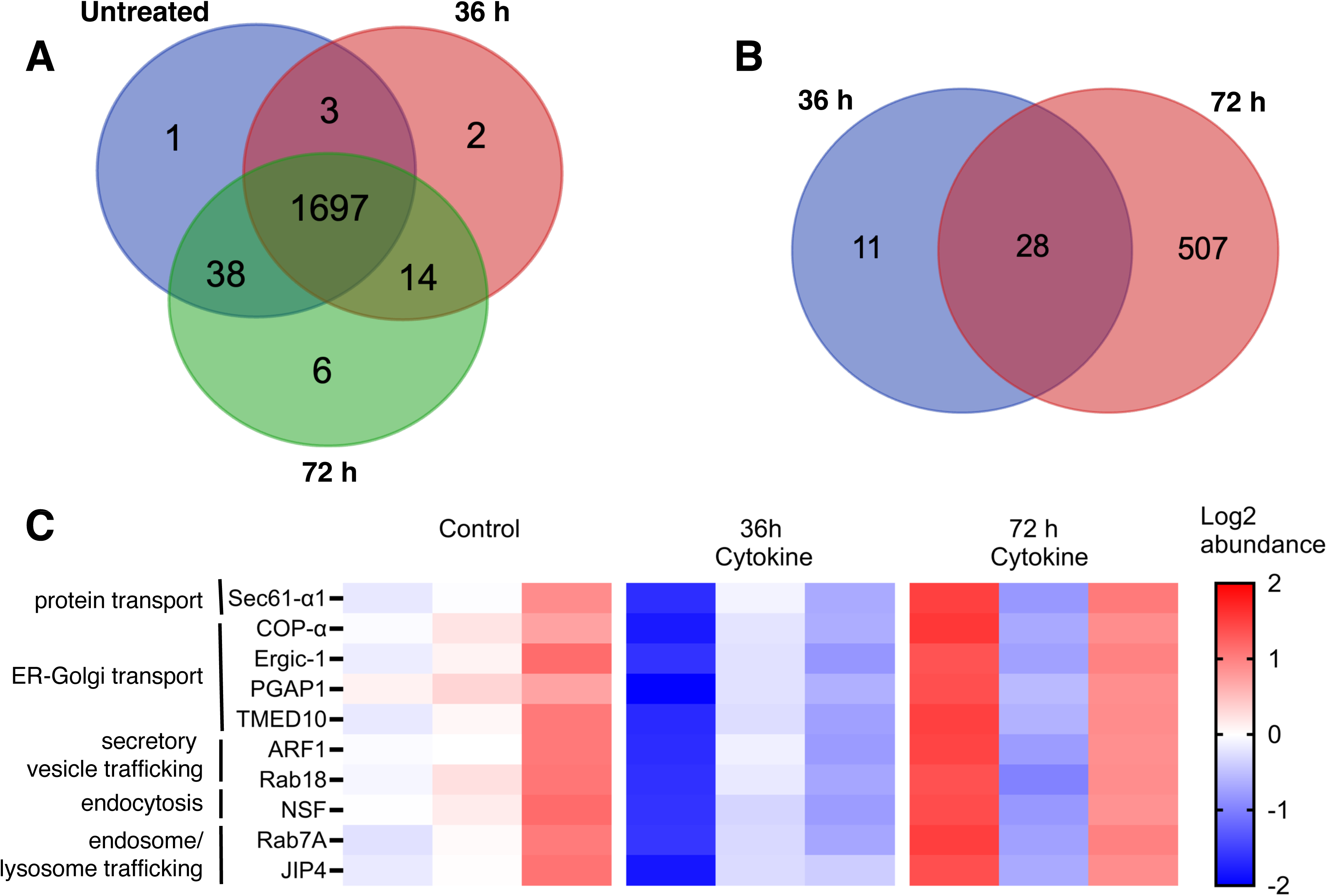
Carbonylated proteins in MIN6 cells treated with proinflammatory cytokines. **A:** Venn diagram showing the number of distinct proteins detected by LC-MS following purification of carbonylated proteins from cells treated with the cytokine cocktail for the indicated time. Only proteins from which ≥ 4 peptides were detected are included. **B:** Venn diagram showing the number of distinct proteins that were ≥1.5-fold enriched in the treatment samples relative to controls. **C:** Heat map showing relative abundance of PC for representative selected proteins. Full lists are given in the Supporting Information, **File S3**.

Proteomic results suggest that patterns of PC induced by proinflammatory cytokines partially overlap with those induced by direct 4-HNE application (**Fig. S3**). GO enrichment analysis showed some pathways in common with both NOD islets and 4-HNE-treated cells, most notably intracellular protein transport (including vesicle trafficking) (**Tables 1-3**). However, the cellular compartments targeted by cytokine-induced protein carbonylation were notably enriched in mitochondria and endoplasmic reticulum proteins, consistent with induction of endogenous ROS production from these compartments (**Table 3**). Correspondingly, the ER-dominant pathways of protein folding and antigen presentation were also among the top 10 enriched biological processes in the cytokine-treated cells. These pathways were also observed among the top ten enriched pathways in NOD islets (**Table 1**) but not 4-HNE treated cells (**Table 2**). Overall, cytokine and 4-HNE treatments led to distinct but overlapping patterns of protein carbonylation in MIN6 cells, and the pattern observed in NOD islets is a composite of those induced by these two physiological sources of oxidative stress. In all three stress conditions, membrane trafficking proteins are abundantly represented among the identified PC targets.

**Table 3:** Gene Ontology categories among carbonylated proteins 1.5-fold enriched following 72-hour cytokine treatment.

| Biological Process category | Log FDR | Cellular Compartment category | Log FDR |
| --- | --- | --- | --- |
| intracellular protein transport | -15.5 | mitochondrion | -52.8 |
| endoplasmic reticulum to Golgi vesicle-mediated transport | -10.0 | cytosol | -41.2 |
| chromatin looping | -9.7 | endoplasmic reticulum | -30.0 |
| cytoplasmic translation | -9.5 | endoplasmic reticulum membrane | -29.8 |
| protein transport | -8.9 | cytoplasm | -26.0 |
| translation | -7.6 | mitochondrial matrix | -17.3 |
| protein folding | -7.2 | mitochondrial inner membrane | -16.4 |
| response to virus | -6.9 | ribosome | -14.8 |
| fatty acid beta-oxidation | -6.8 | protein-containing complex | -13.1 |
| antigen processing and presentation | -6.7 | Golgi apparatus | -12.7 |

## Discussion

### Summary of key findings

Inflammation of pancreatic islets during pre-T1D generates oxidative stress, which is a key contributor to the dysfunctional insulin secretion that precedes β-cell destruction. Increased concentrations of ROS induce lipid peroxidation of unsaturated fatty acids, which in turn produces reactive aldehydes that carbonylate proteins. In the present study, we have combined proteomic identification of carbonylated proteins with measurements of insulin secretion to make the following significant observations: (i) over 1000 proteins are targets of PC in pre-diabetic NOD mouse islets and cultured β-cells; (ii) islets from NOD mice and β-cells treated with cytokines or 4-HNE showed an increase in PC compared to their control counterparts; (iii) GO analysis revealed that all tested stress conditions led to increased carbonylation of proteins associated with processes central to β-cell function including membrane trafficking; (iv) consistent with this finding, PC impairs insulin secretion by ∼50% in cultured MIN6 and INS-1 GRINCH cells incubated with cytokines or 4-HNE compared to untreated cells; (v) surprisingly, 4 HNE exposure inhibited insulin secretion within 5-10 minutes. This timescale is too fast to be explained by changes in gene expression, suggesting that PC of secretory or signaling proteins may directly impact insulin release.

### Possible origins of rapid inhibition of insulin secretion

Increased PC and inhibition of insulin secretion were both observed within 5-10 minutes of 4-HNE addition to cultured INS-1 (GRINCH) and MIN6 β-cells (**Figs. 2 and 3**). Previous studies have ascribed impaired insulin secretion in diabetes to a range of factors including decreased insulin expression, decreased granule acidification, and β-cell dedifferentiation (55). While these processes are clearly important in long-term disease, their timescales suggest they are unlikely to account for the rapid inhibition upon 4-HNE exposure. Rather, we speculate that rapid inactivation via PC may impair the function of one or more required proteins. Although we cannot exclude the possibilities that 4-HNE may activate rapid signaling pathways or directly impair Ca^2+^ influx on the minutes timescale, we note that Ca^2+^ signaling was not among the top cellular pathways identified in any of our GO analyses (**Tables 1-3**).

Across all stress conditions tested, a significant increase in PC was observed in proteins associated with secretory vesicle trafficking, exocytosis, and endocytosis, including 27 different Rab GTPases, the Rab effector Slp-4/granuphilin, N-ethylmaleimide sensitive factor (NSF), adaptor protein 2 (AP2) complex, and clathrin (**Table S1**). Together, the carbonylated proteins comprise most of the members of the KEGG pathway of synaptic vesicle exo/endocytosis (56), a process highly similar to insulin release (**Fig. 7A**). Many of these proteins are required for normal insulin secretion in β-cells (57–61), and several including Rab1a, Rab6a, and AP2-μ have been associated with diabetes in genome-wide association studies (62). Rab GTPases were also identified among key targets of PC in cholestatic liver disease (63), suggesting that this key class of membrane trafficking proteins may be generally sensitive to PC. We also note that insulin and/or proinsulin had increased carbonylation in the NOD and 4HNE experiments (**Table S1**). Further work is needed to clarify how carbonylation of individual proteins contributes to the inhibition of insulin secretion. Because we observe many of the same proteins to be carbonylated in 4-HNE treated cells and NOD islets, this process could represent a novel mode of inhibited insulin secretion in early T1D.

**Figure 7:**
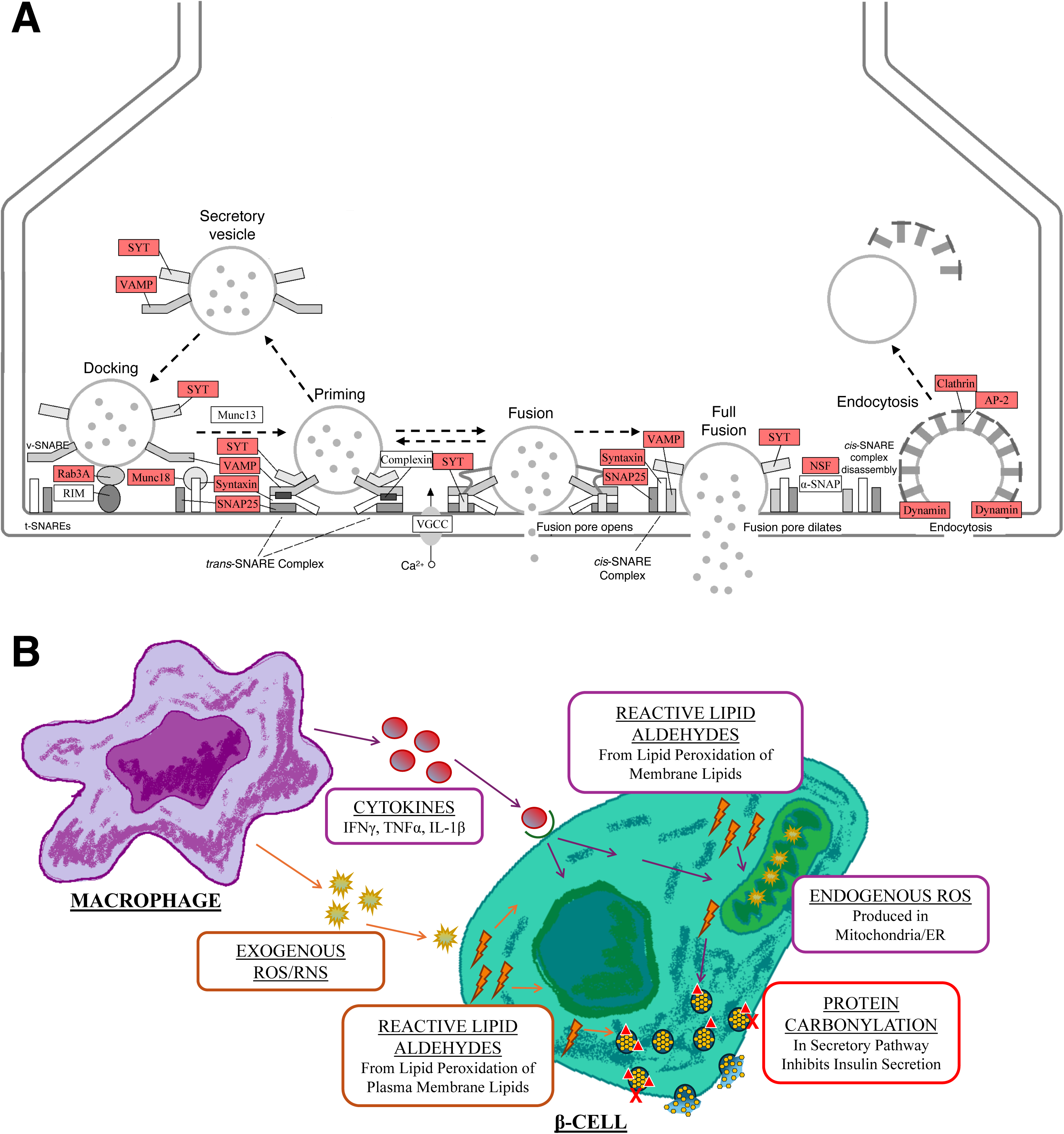
Protein carbonylation and insulin secretion in pre-T1D. **A:** Carbonylated proteins in exocytosis and endocytosis. Shown is a modified KEGG pathway diagram of the synaptic vesicle cycle (https://www.kegg.jp/pathway/map04721, reproduced with permission (56)), which involves the same families of proteins as insulin secretion at this level of detail. Proteins highlighted red were identified as enriched in one or more of the experiments described in this paper (**Table S1**). **B:** Cellular model of PC in T1D. Islet inflammation leads to both extracellular (exogenous, orange arrows) and intracellular (endogenous, purple arrows) sources of ROS, which lead to carbonylation of distinct but overlapping sets of proteins and both inhibit insulin secretion by ∼50%. Original artwork by K.R. Schultz.

### PC pattern in NOD mouse islets is consistent with a combination of ROS sources in pre-T1D

Our findings support roles for exogenous and endogenous sources of ROS and their subsequent production of toxic lipid aldehydes in pre-T1D islets. The data reveal carbonylation of a wide range of proteins in pre-T1D NOD mouse islets (**Fig. 1 and File S1**), most of which likely reflect changes in β-cells. We note that β-cells comprise ∼80% of healthy mouse islets and somewhat less in NOD mice experiencing insulitis (64; 65). Indeed, the 94 carbonylated proteins that were exclusively detected in NOD but not WT islets (**Figure 1C**) cluster with GO terms for antigen presentation and lamellipodia formation, suggesting that these proteins may have arisen from infiltrating immune cells (**Table S3)**. Therefore, to verify PC occurring in β-cells under inflammatory stress conditions, we used the MIN6 β-cell line treated with 4-HNE or proinflammatory cytokines.

Comparing PC patterns in these model systems reveals insights into the β-cell pathways targeted. Pathways of secretory protein trafficking, ribosomal translation, gene regulation, and metabolism are targets of PC under all three conditions (**Tables 1-3**). GO analysis of carbonylated proteins in 4-HNE treated MIN6 cells replicated most of the biological processes associated with PC in the NOD islets, including protein transport, membrane trafficking, and gene regulation (**Tables 1-2 and Fig. S3**). Proinflammatory cytokines also effected carbonylation of proteins in many of these processes as well as proteins that typically localize to mitochondria and ER, which is not surprising as these are two major sources of ROS in β-cells (11; 66). On the other hand, 589 proteins had increased carbonylation in NOD islets and 4-HNE treated cells but not cytokine-treated cells (**Fig. S3**). As a whole, these results suggest that protein carbonylation in pre-diabetic β-cells likely arises from a combination of sources including those induced by cytokine signaling in the β-cell (endogenous) and those derived from compounds released by inflammatory cells (exogenous) (**Fig. 7B**). Further studies are needed to clarify how these sources together contribute to the progression of T1D.

## List of abbreviations

4-HNE: 4-Hydroxynonenal
AP2: Adaptor Protein 2
ATP: Adenosine Triphosphate
BCA: Bicinchoninic Acid
BSA: Bovine Serum Albumin
COPII: Coat Protein Complex II
DMEM: Dulbecco’s Modified Eagle Medium
ELISA: Enzyme-Linked Immunosorbent Assay
ER: Endoplasmic Reticulum
FBS: Fetal Bovine Serum
Gclc: Glutamate-Cysteine Ligase Catalyzed Subunit
GFP: Green Fluorescent Protein
GO: Gene Ontology
GRINCH: Glucose-Responsive Insulin-Secreting C-Peptide-Modified Human-Proinsulin
GPX1: Glutathione Peroxidase
HEPES: 4-(2-hydroxyethyl)-1-piperazineethanesulfonic acid
Hprt: Hypoxanthine Guanine Phosphoribosyltransferase
IFNγ: Interferon-γ
IL-1β: Interleukin-1β
LC-MS: Liquid Chromatography – Mass Spectrometry
MIN6: Mouse Insulinoma 6
MS: Mass Spectrometry
NOD: Non-Obese Diabetic
NOX2: NADPH oxidase 2
Nqo1: NAD(P)H Quinone Dehydrogenase 1
NSF: N-ethylmaleimide Sensitive Factor
P4Hb: Protein Disulfide Isomerase
PBS: Phosphate Buffered Saline
PC: Protein Carbonylation
PCR: Polymerase Chain Reaction
PTM: Post-Translational Modification
qPCR: Quantitative Polymerase Chain Reaction
qRT-PCR: Quantitative Reverse Transcription Polymerase Chain Reaction
RNS: Reactive Nitrogen Species
ROS: Reactive Oxygen Species
SEM: Standard Error of the Mean
sfGFP: Superfolder Green Fluorescent Protein
SLP: Synaptotagmin-Like Protein
SOD2: Superoxide Dismutase 2
STX5: Syntaxin-5
TIRFM: Total Internal Reflection Fluorescence Microscopy
TNFα: Tumor Necrosis Factor-α
WT: Wild Type

## Acknowledgments

This work was supported by a pilot/feasibility award to J.D.K. through NIDDK grant # P30-DK116073. I.L. and V.S.B. were also supported through awards from the Office of Undergraduate Research at the University of Colorado Denver. INS-1 (GRINCH) cells were a gift from Peter Arvan, and MIN6 cells were a gift from the late John Hutton. We thank Scott Beard at the Barbara Davis Center islet core facility for expert islet extraction and Cole Michel at the Skaggs School of Pharmacy mass spectrometry core facility for expert proteomics data collection and analysis. We thank Arun Anantharam for training on TIRFM-based secretion measurements. We also thank Lori Sussel, Kathryn Haskins, Jane Reusch, Paul Rozance, Nicole Farnsworth, and Richard Benninger for critical discussions.

