## Supporting Information for "Inflammatory Oxidative Stress Compounds Inhibit Insulin Secretion through Rapid Protein Carbonylation"

### Supplementary Tables and Figures for Inflammatory Oxidative Stress Compounds Inhibit Insulin Secretion through Rapid Protein Carbonylation

Emma F. Saunders[1†], Katherine R. Schultz[1†], Isaiah Lowe[1], Aimee L. Anderson[2], Vrishank S. Bikkumalla[1], David Soto[1], Nhi Y. Tran[1], Sharon Baumel-Alterzon[3], Jefferson D. Knight[1*], and Colin T. Shearn[2*]

[1]Department of Chemistry, University of Colorado Denver, USA
[2]Department of Pediatrics, University of Colorado Anschutz Medical Campus, Aurora, USA
[3]Department of Molecular and Cellular Endocrinology, Arthur Riggs Diabetes & Metabolism Research Institute at City of Hope, Duarte, CA, USA

†Equal contributions

**Table S1: Carbonylated exocytosis/endocytosis proteins identified in this study.** Check marks indicate that carbonylation was enriched >2-fold (NOD, 4HNE) or >1.5-fold (cytokine) in the indicated experiments. See **Supplementary Files 1-3** for full protein lists.

| **Protein** | **NOD** | **4HNE** | **Cytokine** | | **Protein** | **NOD** | **4HNE** | **Cytokine** |
|---|---|---|---|---|---|---|---|---|
| Rab1A | ✓ | ✓ | ✓ | | INS1 | | ✓ | |
| Rab1B | ✓ | ✓ | ✓ | | INS2 | ✓ | ✓ | |
| Rab2A | ✓ | ✓ | | | **Syts and Syt-like** | | | |
| Rab2B | | ✓ | | | Syt-2 | | ✓ | |
| Rab3A | ✓ | ✓ | | | SytL4 (granuphilin) | ✓ | ✓ | |
| Rab3B | ✓ | ✓ | ✓ | | SytL5 | | ✓ | |
| Rab3C | | | ✓ | | **SNARE complex** | | | |
| Rab3D | ✓ | ✓ | | | Syntaxin-5 | | ✓ | |
| Rab4B | ✓ | | | | Syntaxin-7 | | ✓ | |
| Rab5A | ✓ | ✓ | | | Syntaxin-12 | | ✓ | |
| Rab5B | ✓ | | ✓ | | SNAP25 | | ✓ | |
| Rab5C | ✓ | ✓ | | | SNAP47 | | ✓ | |
| Rab6A | ✓ | ✓ | ✓ | | VAMP4 | | ✓ | |
| Rab7A | ✓ | ✓ | ✓ | | **Syntaxin-binding** | | | |
| Rab8A | ✓ | ✓ | ✓ | | Stxbp1 (Munc18-1) | ✓ | ✓ | |
| Rab9B | | ✓ | | | Stxb5 (Tomosyn-1) | | ✓ | |
| Rab10 | ✓ | | | | Stxb5L (Tomosyn-2) | | ✓ | |
| Rab11B | ✓ | ✓ | | | **Endocytosis** | | | |
| Rab12 | | ✓ | ✓ | | NSF | ✓ | ✓ | ✓ |
| Rab14 | ✓ | ✓ | | | Clathrin heavy chain | ✓ | ✓ | |
| Rab18 | ✓ | ✓ | ✓ | | AP2-α1 subunit | ✓ | ✓ | |
| Rab21 | ✓ | ✓ | ✓ | | AP2-α2 subunit | ✓ | ✓ | |
| Rab27A | ✓ | ✓ | | | AP2-β subunit | ✓ | ✓ | |
| Rab34 | | ✓ | | | AP2-μ subunit | ✓ | ✓ | |
| Rab35 | | | ✓ | | AP2-σ subunit | | ✓ | |
| Rab37 | ✓ | ✓ | | | Dynamin-1 | | ✓ | |
| Rab39B | | ✓ | | | Dynamin-2 | | ✓ | |

**Table S2: GO categories among carbonylated proteins >2-fold enriched following 5-minute 4-HNE treatment**

| Biological Process category | Log FDR | Cellular Compartment category | Log FDR |
|---|---|---|---|
| cytoplasmic translation | -48.8 | cytosol | -266.7 |
| protein transport | -47.2 | cytoplasm | -222.6 |
| intracellular protein transport | -42.9 | nucleus | -155.9 |
| translation | -38.6 | nucleoplasm | -119.9 |
| translation at postsynapse | -38.6 | ribonucleoprotein complex | -79.2 |
| mRNA processing | -38.3 | postsynapse | -78.1 |
| translation at presynapse | -37.7 | protein-containing complex | -68.5 |
| chromatin remodeling | -31.3 | synapse | -65.7 |
| cell division | -29.7 | presynapse | -57.7 |
| protein import into nucleus | -27.3 | glutamatergic synapse | -54.9 |

**Table S3: GO analysis of 94 carbonylated proteins detected in NOD but not WT islets**

| Biological Process category | Log FDR | Cellular Compartment category | Log FDR |
|---|---|---|---|
| cellular response to type II interferon | -1.8 | Cytoplasm | -9.6 |
| cellular response to interferon-beta | -1.5 | Cytosol | -8.1 |
| innate immune response | -1.3 | Stress fiber | -4.0 |
| antigen processing and presentation | -1.3 | Lamellipodium | -3.5 |
| | | Protein-containing complex | -3.4 |
| | | Cytoskeleton | -3.3 |
| | | Perinuclear region of cytoplasm | -3.2 |
| | | Nucleus | -3.2 |
| | | Actin cytoskeleton | -3.1 |
| | | Mitochondrion | -2.3 |

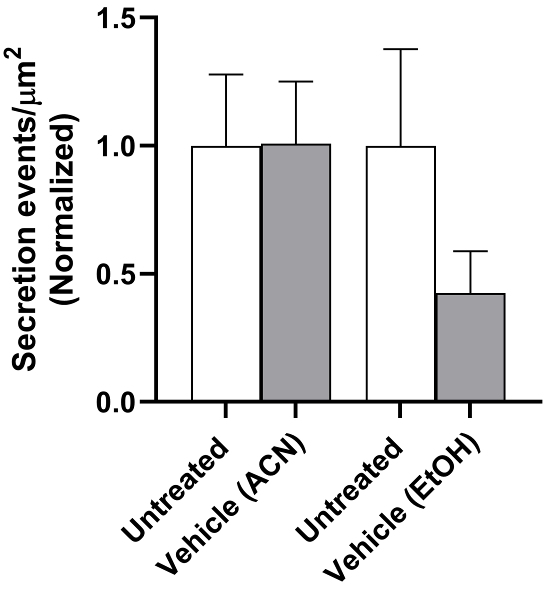


**Figure S1: Vehicle controls for TIRF secretion assay.** INS-1 (GRINCH) cells were stimulated with 60 mM KCl for 1 min, before or after the addition of solvent vehicle [acetonitrile (ACN) or ethanol (EtOH)] in an equivalent volume to that used to deliver 100 uM 4-HNE. The number of secretion events in each sample was counted and normalized to the imaged cell area. Acetonitrile was used as the solvent for all 4-HNE experiments in the main paper.

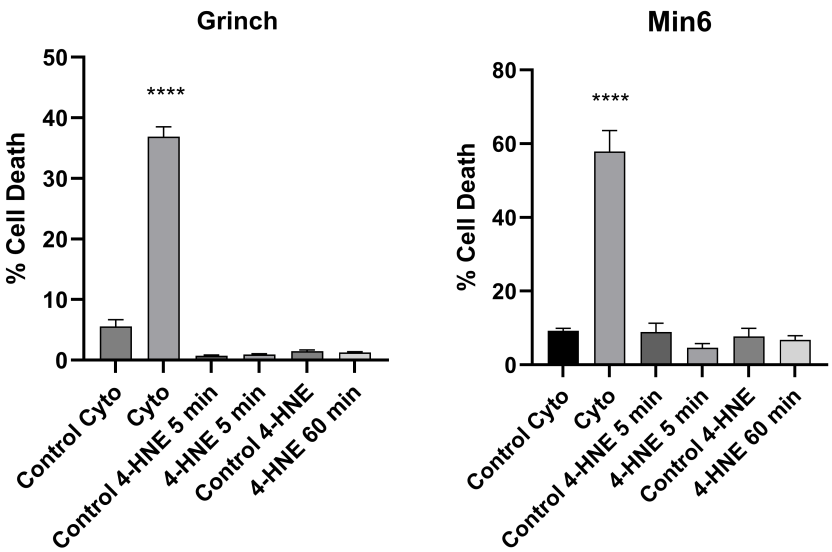


**Figure S2: Effects of cytokine (Cyto) and 4-HNE treatment on cell death**. INS-1 (GRINCH) or MIN6 cells were treated with the cytokine cocktail for 72h or with 100 μM 4-HNE for 5 min or 1h. Cell death was measured using DAPI as described in Methods. Data shown are mean ± SEM from at least 4 replicate measurements each. ****$p<0.001$; n.s. not significant.

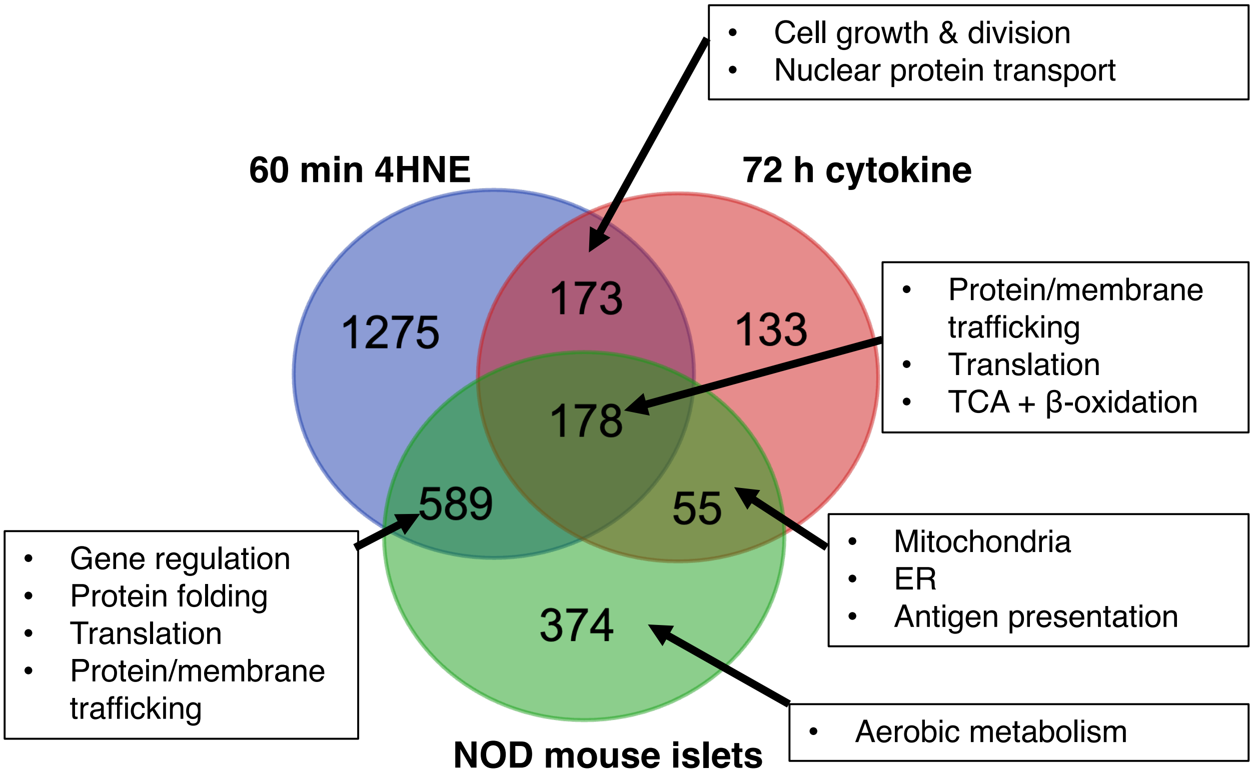


**Figure S3: Venn diagram showing overlap of carbonylated proteins identified** in NOD mouse islets (green, >2-fold enriched vs. WT control), 4HNE-treated MIN6 cells (blue, 60 min treatment, proteins >2-fold enriched vs. untreated cells), and cytokine-treated MIN6 cells (red, 72-h treatment, >1.5-fold enriched vs. untreated cells). Boxes highlight key GO pathways and cellular compartments identified from DAVID analysis of the proteins in each indicated shaded region. The most significant category of carbonylated proteins identified in the NOD islets but not the MIN6 cells was mitochondrial respiration, likely reflecting a lower abundance of these proteins in insulinoma-derived cell lines due to their Warburg-like metabolism. Conversely, pathways represented by carbonylated proteins enriched in the insulinoma-derived cells but not NOD islets included cell division and chromatin remodeling; this is not surprising, as these proteins are more abundant in rapidly dividing cells. Despite these expected limitations, 69% of the proteins with increased carbonylation in NOD mouse islets were also identified in MIN6 cells treated with 4-HNE and/or cytokines.  Full listings of proteins and GO pathways identified in this analysis are in **SI File S4**.
